# Inflammatory pain in mice induces light cycle-dependent effects on sleep architecture

**DOI:** 10.1101/2024.08.28.610124

**Authors:** Dominika J. Burek, Khairunisa Mohamad Ibrahim, Andrew G. Hall, Ashish Sharma, Erik S. Musiek, Jose A. Morón, William A. Carlezon

**Affiliations:** Basic Neuroscience Division, McLean Hospital, Belmont, MA, 02478, USA; Department of Psychiatry, Harvard Medical School, Boston, MA, 02115, USA; Department of Anesthesiology, Washington University Pain Center, St. Louis, MO, 63110, USA; Washington University in St. Louis, School of Medicine, St. Louis, MO, 63110, USA; Department of Neuroscience, Washington University in St. Louis, St. Louis, MO, 63110, USA; Department of Psychiatry, Washington University in St. Louis, St. Louis, MO, 63110, USA; Department of Neurology, Washington University School of Medicine, St Louis, Missouri, 63110, USA; Center on Biological Rhythms and Sleep (COBRAS), Washington University School of Medicine, St. Louis, Missouri, 63110, USA

**Author notes:** Co-corresponding authors: correspondence should be sent to and. Co-first authors (Dominika J. Burek, Khairunisa Mohamad Ibrahim).

## Abstract

As a syndrome, chronic pain comprises physical, emotional, and cognitive symptoms such as disability, negative affect, feelings of stress, and fatigue. A rodent model of long-term inflammatory pain, induced by complete Freund’s adjuvant (CFA) injection, has previously been shown to cause anhedonia and dysregulated naturalistic behaviors, in a manner similar to animal models of stress. We examined whether this extended to alterations in circadian rhythms and sleep, such as those induced by chronic social defeat stress, using actigraphy and wireless EEG. CFA-induced inflammatory pain profoundly altered sleep architecture in male and female mice. Injection of the hind paw, whether with CFA or saline, reduced some measures of circadian rhythmicity such as variance, period, and amplitude. CFA increased sleep duration primarily in the dark phase, while sleep bout length was decreased in the light and increased in the dark phase. Additionally, CFA reduced wake bout length, especially during the dark phase. Increases in REM and SWS duration and bouts were most significant in the dark phase, regardless of whether CFA had been injected at its onset or 12 hours prior. Taken together, these results indicate that inflammatory pain acutely promotes but also fragments sleep.

## INTRODUCTION

An estimated 20 to 40% of Americans live with chronic pain, half of whom report daily suffering^1,2^. Chronic pain can be the result of a nerve injury or inflammation, a symptom of other disorders and syndromes, or idiopathic (unknown cause)^3^. Regardless of the cause, chronic pain has profound effects on daily life: patients report their pain interferes with independence, mobility, work, relationships, and sleep quality^4,5^. Fragmented sleep and debilitating fatigue are especially burdensome for those with painful autoimmune disorders^6^.

Considering its disruptive effects, pain can be conceptualized as a persistent stressor. Indeed, the emotional toll of feelings like catastrophizing, helplessness, and despair perpetuates cycles of stress and pain^7,8^. Approximately 30% of chronic pain patients meet diagnostic criteria for psychiatric disorders commonly associated with stress, such as Major Depressive Disorder (MDD)^9,10^. Sleep dysregulation is a prominent feature of depressive disorders, and includes reduced latency to rapid-eye-movement (REM) sleep and increased REM duration, which are well-characterized biomarkers of MDD^11,12^. We have previously shown that a mouse model used to study traumatic psychosocial stress, chronic social defeat stress (CSDS), promotes both REM and non-REM, or slow-wave, sleep (SWS)^11^. The same CSDS regimen produces anhedonia—a key diagnostic feature of MDD—that develops over a virtually identical time course^13^. Linking these signs mechanistically, we showed that neural manipulations that regulate motivation also regulate sleep patterns^14^. Even acute stressors such as a single instance of social conflict or restraint have also been shown to increase SWS sleep, in both humans and rodents^15–17^.

This overlap among pain, stress, and sleep dysregulation may be attributed to how these experiences engage and are moderated by the same circuitry, such as the hypothalamic-pituitary-adrenal (HPA) axis, mesolimbic reward pathway, and lateral hypothalamus^14,18–20^. Another factor that may contribute to this overlap, particularly in the case of autoimmune or inflammatory pain, is the immune system^21^. Pro-inflammatory signaling molecules such as the cytokines IL-1β, IL-6, and TNF-α are circadian-regulated and bidirectionally modulate sleep in both illness and healthy conditions^6,21^. Cytokines are also stimulated by CSDS and inflammatory pain, raising the possibility of shared mechanisms^22–25^.

The present studies were designed to characterize the effects of inflammatory pain on sleep. We hypothesized that inflammatory pain would produce the same effects on sleep as stress^11,14^. To model chronic inflammatory pain, we used Complete Freund’s Adjuvant (CFA)-induced inflammation, a well-characterized rodent model of long-term and persistent pain^26,27^. CFA comprises mycobacterium antigen, which evokes inflammation both locally at the site of injection and throughout the central nervous system^24,28,29^. When injected into the hind-paw, CFA causes swelling, erythema, and putatively painful thermal and mechanical hypersensitivity that can last for six weeks^30^. Although there are dozens of reports describing the behavioral phenotype of CFA-induced inflammatory pain, our preclinical meta-analysis indicated that its effects on most classical assays of anxiety- and depressive-like behaviors are small and heterogeneous^31^. Interestingly, the most prominent effects are observed in more naturalistic behaviors such as wheel running, nesting, burrowing, and in translationally-relevant signs such as anhedonia^18,32–35^. Since classical phenotyping assays have not yet identified strong biomarkers that have enabled development of novel therapeutics, more objective and continuous measures may provide a more complete understanding of how pain acts as a stressor^36^. To better understand the nuanced ways in which CFA-induced inflammatory pain affects sleep, we leveraged locomotor activity tracking, piezoelectric sensors, and EEG recordings. Our studies reveal that although CFA has little effect on circadian rhythms, it strongly promotes sleep, causing suppression of wakefulness and reductions in wake bout length and corresponding increases in both duration and bouts of both REM and SWS. Furthermore, we discovered that CFA promotes sleep most prominently in the dark phase of the light cycle, when mice are normally more likely to be active.

## METHODS

### Animals

Male and female C57BL/6J mice, 7 weeks old, were obtained from Jackson Laboratories (Bar Harbor, ME, USA) and acclimated to the vivarium for one week. Mice were housed in temperature- (21 ± 2°C) and humidity-(50 ± 20%) controlled vivarium rooms with food, water, and nesting material available *ad libitum*. All procedures and methods were approved by the Washington University in St. Louis Institutional Animal Care and Use Committee or the McLean Hospital Institutional Animal Care and Use Committee, and performed in accordance with the National Institutes of Health’s (NIH) Guide for the Care and Use of Animals.

### Actigraphy-based quantification of circadian locomotor activity in Light-Dark and Dark-Dark cycle

Mice were individually housed in Ancare N10 mouse cages (Ancare Corp., Bellmore, NY) equipped with passive infrared sensor wireless nodes (Actimetrics, Wilmette, IL). Daily locomotor activity was recorded at 1-minute intervals using the ClockLab Wireless data collection program v4.129 (Actimetrics, Wilmette, IL) under a 12h light/12h dark (LD) cycle (lights on from 0600, zeitgeber time [ZT] 0, to 1800 hours; 300 lux) for 7 days baseline, followed by 7 days post-saline injection, and lastly 7 days post-CFA injection. In Dark-Dark circadian locomotor activity studies, activity was recorded daily under 12h light/12h dark (LD) cycle (lights on from 0600 [ZT 0] to 1800 hours [ZT 12]; 300 lux) for the first 6 days, followed by constant dark (DD) conditions for the next 15 days (1 additional day baseline, 7 days post-saline and 7 days post-CFA injections). Mice had ad libitum access to food, water, and nesting material throughout the experiments. Period length (tau) was determined through χ2 periodogram analysis, rhythm strength was measured using relative fast Fourier transform analysis, and other non-parametric features of circadian locomotor activity were assessed with ClockLab Analysis v6.1.02 (Actimetrics, Wilmette, IL). Routine animal husbandry during the dark phase was conducted under dim red light (< 10 lux). All hind paw injections were done 3 hours after “lights-on” (0900 hours, ZT 3) for the LD cycle or at 0900 hours (ZT 3) for the DD cycle.

### Actigraphy-based quantification of sleep and wake states

To measure sleep-like behavior in mice, we used an automated piezoelectric sleep monitoring system (Adapt-A-Base, Model #MB-ACN10, Signal Solutions, Lexington, KY). This highly sensitive system uses short-time (2 seconds) pressure signal segments to determine sleep/wake states. The non-invasive, high-throughput system detects breathing and gross body movements to analyze unsupervised sleep/wake patterns with 90% accuracy compared to EEG/EMG-based methods^37,38^. When the animals were asleep, the main pressure changes were due to breathing, producing an accurate respiratory trace. Sleep states showed quasi-periodic signals with minimal amplitude variations. In contrast, the wakefulness state exhibited irregular, transient, high-amplitude pressure changes linked to body movements and weight shifting. Even during “quiet rest,” slight head or other movements were enough to differentiate rest from sleep with accuracy comparable to EEG/EMG^38^. Mice were individually housed in Ancare N10 mouse cages (Ancare Corp., Bellmore, NY) and placed on the Adapt-A-Base systems in sound- and light-proof circadian cabinets (Actimetrics, Wilmette, IL). They were kept under a 12h light/12h dark cycle (LD) (lights on from 0600 [ZT 0] to 1800 hours; 300 lux), with ad libitum access to food, water, and nesting material. Each cage contained 160 g of corncob bedding, 220 g of food pellets, and 380 mL of water, ensuring consistent cage weight except for minor variations in mouse body weight. Sleep/wake states were recorded continuously for 15 days using the PiezoSleep v2.08 program (Signal Solutions, Lexington, KY) without any disturbance except for saline or CFA hind paw injections. All hind paw injections were done 3 hours after “lights-on” (0900 hrs, ZT 3). Sleep data were analyzed for various sleep features of individual mice using SleepStats v2.18 (Signal Solutions, Lexington, KY). Sleep or wake bout lengths were scored using 30-second epochs.

### EEG- and EMG-based quantification of sleep telemetry: implant surgeries

Mice were implanted with HD-X02 wireless telemetry devices from Harvard Bioscience, Inc. dba Data Sciences International (DSI; St. Paul, MN, USA) to enable continuous recording of electroencephalograms (EEG), electromyogram (EMG), locomotor activity, and subcutaneous temperature. Surgical procedure followed DSI-provided instructions. Mice were anesthetized with 2% isoflurane and immobilized on a stereotaxic instrument. A small incision was made from the medial skull to posterior neck, through which a subcutaneous pocket on the back was opened using lubricated forceps. Transmitters were inserted lateral to the spine, midway between the fore and hind limbs. Two incisions were made in the trapezius muscle using a 21 G needle, through which two insulated biopotential leads (EMG electrodes) were threaded and secured using 6.0 non-dissolvable silk sutures. EEG biopotential leads were attached to two stainless steel screws, inserted into the skull, and lowered until they made contact with dura (relative to bregma: frontal AP +1.0 mm, ML +1.0 mm; parietal AP−3.0 mm, ML −3.0 mm). Screws and EEG leads were secured with dental cement (Infinity by DenMat, Lompoc, CA, USA) and the incision was closed with 6.0 non-dissolvable silk sutures. Triple antibiotic ointment (topical) and lidocaine (2%, topical) were applied to the incision site. Ketoprofen (5 mg/kg, subcutaneous) was administered immediately after surgery, and once daily for 7 days of recovery and post-operative monitoring with food and hydration supplements.

### EEG- and EMG-based quantification of sleep telemetry: data acquisition & physiological recordings

Mice were single housed in Ancare N10 mouse cages (Ancare Corp., Bellmore, NY), under a standard 12h light/12h dark cycle (lights on from 0900 [ZT 0] to 2100 hours; 300 lux). Cages sat on RPC-1 PhysioTel receiver platforms (DSI), which detect an AM radio signal emitted by the HD-X02 transmitter. Receivers connected to a data exchange matrix that continuously uploaded recording data (sampling rate 500 Hz) to Ponemah Software (DSI). Data were analyzed with Neuroscore Software V3.4 (DSI). Raw EEG signal was quantified into relative spectral power in 10-second epochs with Fast Fourier Transformation (FFT) and a Hamming signal window. Frequency bands were defined as delta (0.5– 4 Hz), theta (4–8 Hz), alpha (8–12 Hz), beta (16–24 Hz), and low gamma (30–50 Hz). Vigilance states of active wakefulness (AW), paradoxical or rapid eye movement (REM) sleep, and slow-wave sleep (SWS) were assigned using an automated Neuroscore algorithm, calibrated to each baseline recording. For any 10-second epoch, AW required more than 20% of EEG power to be above the maximum amplitude recorded during either stage of sleep; REM required largest peak in theta power, theta power to be at least 1.1x that of delta power, and minimum EMG amplitude; SWS required largest peak in delta power, delta power to be on average at a ratio of 0.3 to the power of theta, alpha, beta, and gamma combined, and minimum EMG amplitude. Total duration and uninterrupted bouts of each stage were calculated separately for 12 hours of lights-on and 12 hours of lights-off per 24-hour period. Temperature in Celsius and activity counts in arbitrary units were automatically provided by Neuroscore. Activity was normalized to wake duration by dividing activity counts by wake duration, to account for differences in wake duration across time. Baseline was averaged from 7 days of recording, then used to normalize measures within-subjects for recordings after isoflurane exposure, saline injection, and CFA injection.

### Statistical Analyses

Statistical analyses were performed using Prism 9 (GraphPad, Boston, MA, USA) with significance set to p<0.05. Measures for pooled sexes were assessed for normality with Kolmogorov-Smirnov tests, compared with paired t-tests or nonparametric Wilcoxon matched-pairs signed rank tests (SAL vs. CFA), repeated measures one-way ANOVAs and Tukey’s multiple comparisons or nonparametric Friedman and Dunn’s tests (ISO vs. SAL vs. CFA), and compared to a theoretical mean of 100 in one-sample t-tests or nonparametric Wilcoxon signed rank tests. Measures comparing males and females were compared with repeated measures two-way ANOVAs, Tukey’s and Sidak’s multiple comparisons, and means for each sex individually compared to a theoretical mean of 100 in one-sample t-tests. ROUT testing identified two outliers: one excluded from Fig. 3B, for which CFA increased REM sleep duration during the dark phase to 1839.20% of average baseline; and one excluded from Fig. 3H, for which isoflurane increased SWS duration during the dark phase to 249.23% of average baseline. The inclusion vs. exclusion of these data points does not change the significance of the statistical tests. In longitudinal analyses of CFA’s effect on sleep measures in the dark phase (Figs. S2, S4) cage changes on the last day of each week drastically increased wake duration; these days were excluded in analyses. Additionally, transmitter batteries unexpectedly died for two female mice on CFA days 10 and 11; these missing data points required use of a mixed-effects model ANOVA.

## RESULTS

### CFA does not alter circadian rhythms

We first assessed whether CFA hind paw injection affects circadian rhythms. This was assessed through locomotor activity, which was recorded using passive infrared sensor wireless nodes for 7 days of baseline, 7 days after saline injection, and 7 days after CFA injection (**Fig. 1A**).

**Figure 1.**
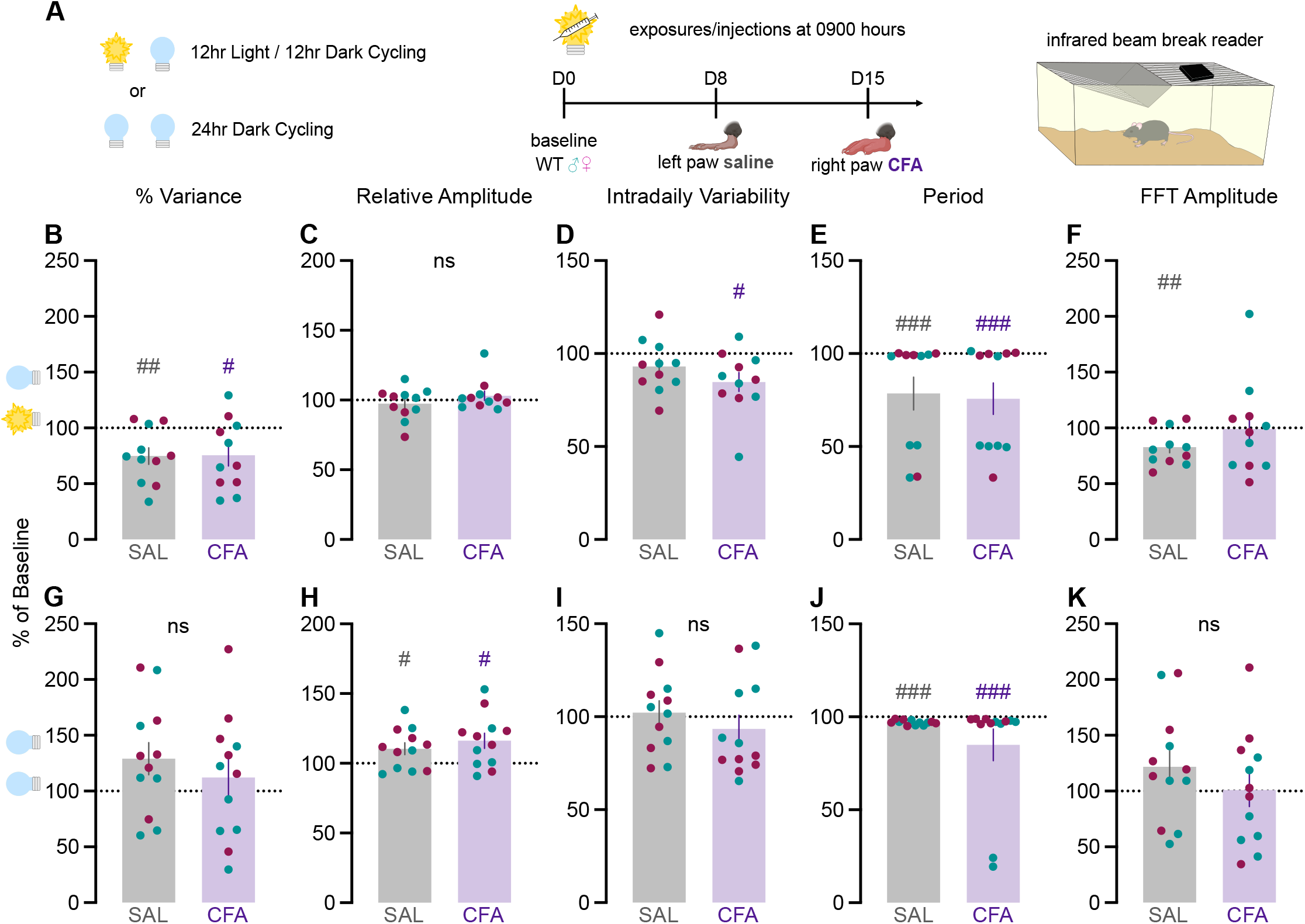
(A) Mice were individually housed in cages equipped with passive infrared sensor wireless nodes. Daily locomotor activity was recorded for 7 days of baseline, 7 days after saline injection, and 7 days after CFA injection. As percent of average baseline (dotted line at 100%), (B) percent variance, (C) relative amplitude, (D) intradaily variability, (E) period, and (F) FFT amplitude under 12 hours light and 12 hours dark cycling conditions. As percent of average baseline, (G) percent variance, (H) relative amplitude, (I) intradaily variability, (J) period, and (K) FFT amplitude under 24 hours constant dark conditions. Teal points indicate males and magenta points indicate females. Ns indicates not significant. # indicates significant difference from theoretical baseline of 100% (# p<0.05, ## p<0.01, ### p<0.001).

Relative to baseline, both saline and CFA reduced percent variance, an indirect measure of daily phase stability (paired t-test t_10_=0.08684, p=0.9325; 1-sample t-test SAL t_10_=3.420, p=0.0066, CFA t_10_=2.595, p=0.0267; **Fig. 1B**). Neither saline nor CFA altered relative amplitude, the ratio of the most active 10 hours to the least active 5 hours (Wilcoxon matched-pairs signed rank p=0.1016; **Fig. 1C**). CFA selectively reduced intra-daily variability, a measure of the fragmentation of a 24-hour rhythm (paired t-test t_10_=2.094, p=0.0627; 1-sample t-test CFA t_10_=3.009, p=0.0131; **Fig. 1D**). For some mice, both saline and CFA significantly reduced the circadian period from an approximate 24-hour baseline (Wilcoxon matched-pairs signed rank test p=0.9658; Wilcoxon signed rank SAL p=0.0010, CFA p=0.0010; **Fig. 1E**). Surprisingly, saline, but not CFA, reduced the highest Fast Fourier Transform (FFT) amplitude estimated by the Lomb-Scargle periodogram (paired t-test t_10_=1.231, p=0.2466; 1-sample t-test SAL t_10_=3.436, p=0.0064, CFA t_10_=0.07744, p=0.9398; **Fig. 1F**). This suggests that CFA injection had no strong effect on circadian rhythm.

We repeated the experiment using a constant darkness protocol to determine if the effect of CFA injection on circadian rhythm was masked by entrainment to the 12h light/12h dark cycle. In the dark-dark protocol, animals were tracked for 6 days of baseline under regular light-dark conditions before switching to constant dark conditions 24 hours prior to saline injection. Neither saline nor CFA had any effect on percent variance (paired t-test t_11_=1.404, p=0.1878; **Fig. 1G**), intradaily variability (paired t t_11_=1.995, p=0.0715; **Fig. 1I**), or FFT amplitude (paired t-test t_11_=1.844, p=0.0923; **Fig. 1K**). Both saline and CFA increased relative amplitude (paired t-test t_11_=2.035, p=0.0667; 1-sample t-test SAL t_11_=2.510, p=0.0290, CFA t_11_=2.954, p=0.0131; **Fig. 1H**) and significantly decreased the period (Wilcoxon matched-pairs signed rank p=0.7695; Wilcoxon signed rank SAL p=0.0005, CFA p=0.0005; **Fig. 1J**).

Together, these findings indicate that CFA did not dramatically alter circadian rhythms during the two weeks of inflammatory pain, regardless of the light cycle.

### CFA increases sleep duration and decreases wake bout length in the first week of inflammation

Although CFA did not impact circadian rhythm, it may still alter sleep quality. To assess this possibility, we examined whether CFA affects sleep quality using an automated piezoelectric sleep monitoring system to detect breathing and gross body movements for analyzing sleep and wake patterns. Mouse activity was recorded for 2 days of baseline, 7 days after saline injection, and 7 days after CFA injection in standard light-dark conditions (**Fig. 2A**). Saline and CFA both increased sleep duration in the light phase relative to baseline, without significant differences across time (RM 1-way ANOVA F_(1.274,19.11)_=1.140, p=0.3152; 1-sample t-test SAL_1-3_ t_15_=4.790, p=0.0002, CFA_1-3_ t_15_=3.819, p=0.0017,CFA_4-6_ t_15_=5.030, p=0.0001; **Fig. 2B**). Sleep bout length in the light phase was significantly reduced over the first three days after CFA injection, relative to saline treatment, the subsequent three days (CFA_4-6_), and to baseline, particularly for females (RM 1-way ANOVA F_(1.604,24.06)_=8.619, p=0.0026, Tukey’s SAL_1-3_ vs. CFA_1-3_ p=0.0064, SAL_1-3_ vs. CFA_4-6_ p=0.8475, CFA_1-3_ vs. CFA_4-6_ p=0.0012; 1-sample t-test CFA_1-3_ t_15_=2.680, p=0.0171; RM 2-way ANOVA: Time F_(1.794,25.12)_=11.00, p=0.0005, Time x Sex F_(2,28)_=5.151, p=0.0125, Tukey’s Female SAL_1-3_ vs CFA_1-3_ p=0.0024, CFA_1-3_ vs CFA_4-6_ p=0.0042; **Fig.2C**). Wake bout length in the light phase was significantly decreased relative to baseline during the three days after saline injection, and even more dramatically reduced during the entire first week after CFA injection, particularly for males (RM 1-way ANOVA F_(1.428,21.42)_=12.06, p=0.0009, Tukey’s SAL_1-3_ vs. CFA_1-3_ p=0.0015, SAL_1-3_ vs. CFA_4-6_ p=0.0382; 1-sample t-test SAL_1-3_ t_15_=2.784, p=0.0139, CFA_1-3_ t_15_=13.05, p<0.0001, CFA_4-6_ t_15_=9.525, p<0.0001; RM 2-way ANOVA Time F_(1.500,21.00)_=13.07, p=0.0005, Tukey’s Male SAL_1-3_ vs. CFA_1-3_ p=0.0131, SAL_1-3_ vs. CFA_4-6_ p=0.0273; **Fig. 2D**).

**Figure 2.**
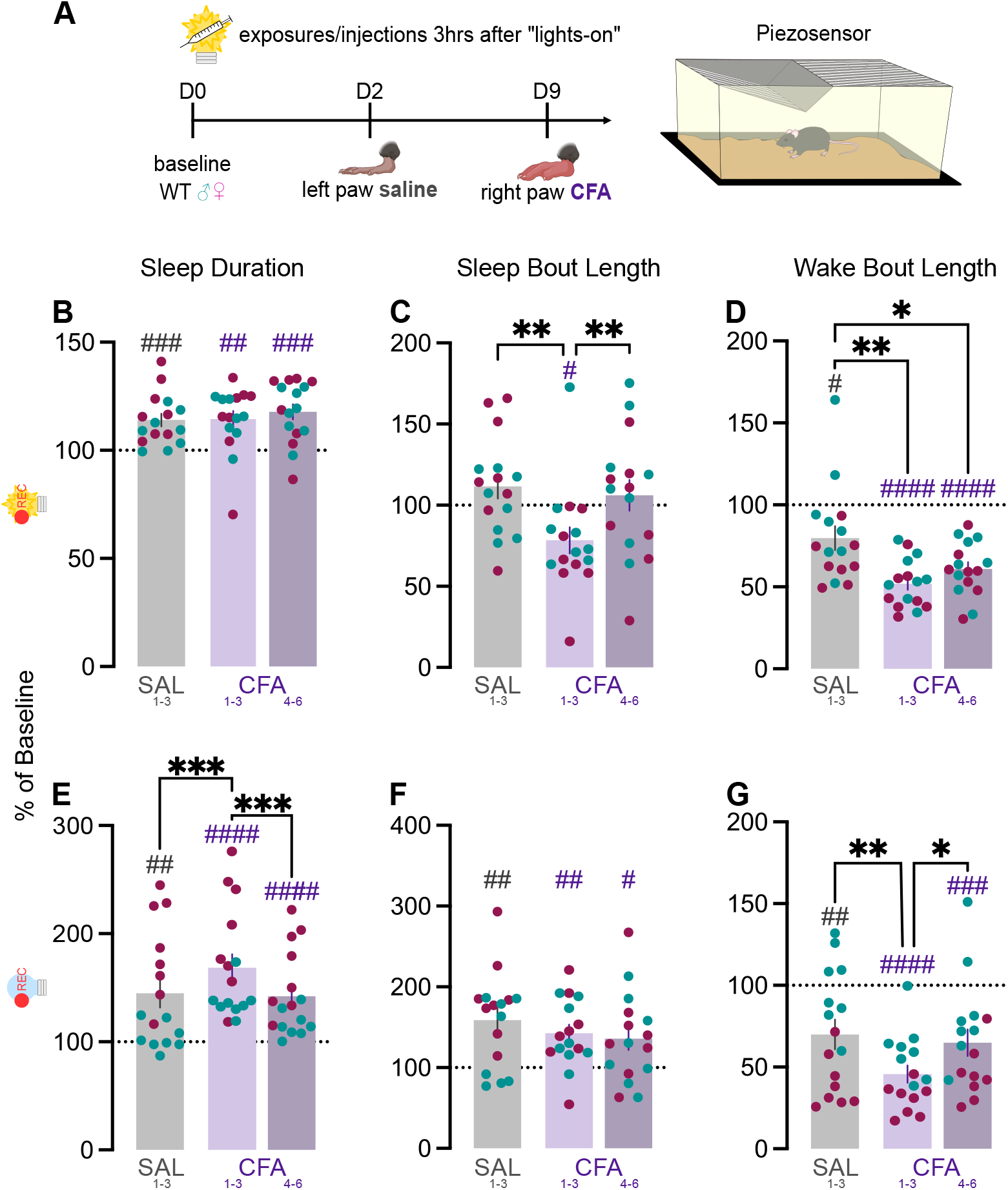
(A) Mice were individually housed in cages on automated piezoelectric sleep monitoring systems. Sleep and wake behavior were recorded for 2 days of baseline, 7 days after saline injection, and 7 days after CFA injection. As percent of average baseline (dotted line at 100%), (B) sleep duration, (C) sleep bout length, and (D) wake bout length during lights-on. As percent of average baseline (E) sleep duration, (F) sleep bout length, and (G) wake bout length during lights-off. Teal points indicate males and magenta points indicate females. Ns indicates not significant. # indicates significant difference from theoretical baseline of 100% (# p<0.05, ## p<0.01, ### p<0.001, #### p<0.0001). * indicate significant group differences (* p<0.05, ** p<0.01, *** p<0.001).

During the dark phase, sleep duration was also increased, with the most pronounced effect in the first three days following CFA injection (CFA_1-3_) (Friedman test F=21.13, p<0.0001; Dunn’s SAL_1-3_ vs. CFA_1-3_ p=0.0003, CFA_1-3_ vs. CFA_4-6_ p=0.0001; Wilcoxon signed rank tests SAL_1-3_ p=0.0034, CFA_1-3_ p<0.0001, CFA_4-6_ p<0.0001; **Fig. 2E**). This was effect was primarily observed in females, though sex differences were apparent across all three time points (RM 2-way ANOVA: Time F_(1.614,22.59)_=21.18, p<0.0001, Sex F_(1,14)_=15.59, p=0.0015, Time x Sex F_(2,28)_=5.699, p=0.0084; Sidak’s Male vs. Female SAL_1-3_ p=0.0038, CFA_1-3_ p=0.0399, CFA_3-6_ p=0.0215; **Fig. 2E)**.

Saline and CFA both increased dark phase sleep bout length relative to baseline, without significant differences across time (RM 1-way ANOVA F_(1.237,18.56)_=1.191, p=0.3020; 1-sample t-test SAL_1-3_ t_15_=4.021, p=0.0011, CFA_1-3_ t_15_=3.998, p=0.0012, CFA_4-6_ t_15_=2.588, p=0.0206; **Fig. 2F**). Wake bout length in the dark phase was significantly reduced over the first three days after CFA injection, relative to saline treatment, the subsequent three days (CFA_4-6_), and to baseline (RM 1-way ANOVA F_(1.797,26.95)_=10.08, p=0.0008, Tukey’s SAL_1-3_ vs. CFA_1-3_ p=0.0033, CFA_1-3_ vs. CFA_4-6_ p=0.0192; 1-sample t-test SAL_1-3_ t_15_=3.327, p=0.0046, CFA_1-3_ t_15_=10.10, p<0.0001, CFA_4-6_ t_15_=4.299, p=0.0006; **Fig.2G**). However, sex differences were observed across all treatment time points, and the reduction in wake bouts was specifically in males (RM 2-way ANOVA Time F_(1.598,22.37)_=11.81, p=0.0006, Sex F_(1,14)_=23.86, p=0.0002, Time x Sex F_(2,28)_=3.582, p=0.0412, Sidak’s Male vs. Female: SAL_1-3_ p=0.0003, CFA_1-3_ p=0.0053, CFA_4-6_ p=0.0451, Tukey’s Male SAL_1-3_ vs. CFA_1-3_ p=0.0098; **Fig. 2G**).

Together, these findings indicate that CFA-induced inflammation increased sleep duration and reduced wake bout length, primarily in the first three nights after injection.

### CFA promotes increased but fragmented REM and SWS

To further characterize the effect of CFA on sleep, we investigated whether the increased sleep was attributable to REM, SWS, or both using wireless telemetry transmitters to enable untethered and continuous recording of EEG, EMG, locomotor activity, and subcutaneous temperature (**Fig. 3A**). EEG and EMG were recorded for 7 days of baseline, 7 days after isoflurane exposure (to control for the anesthesia required to inject implanted animals), 7 days after saline injection, and 21 days after CFA injection.

**Figure 3.**
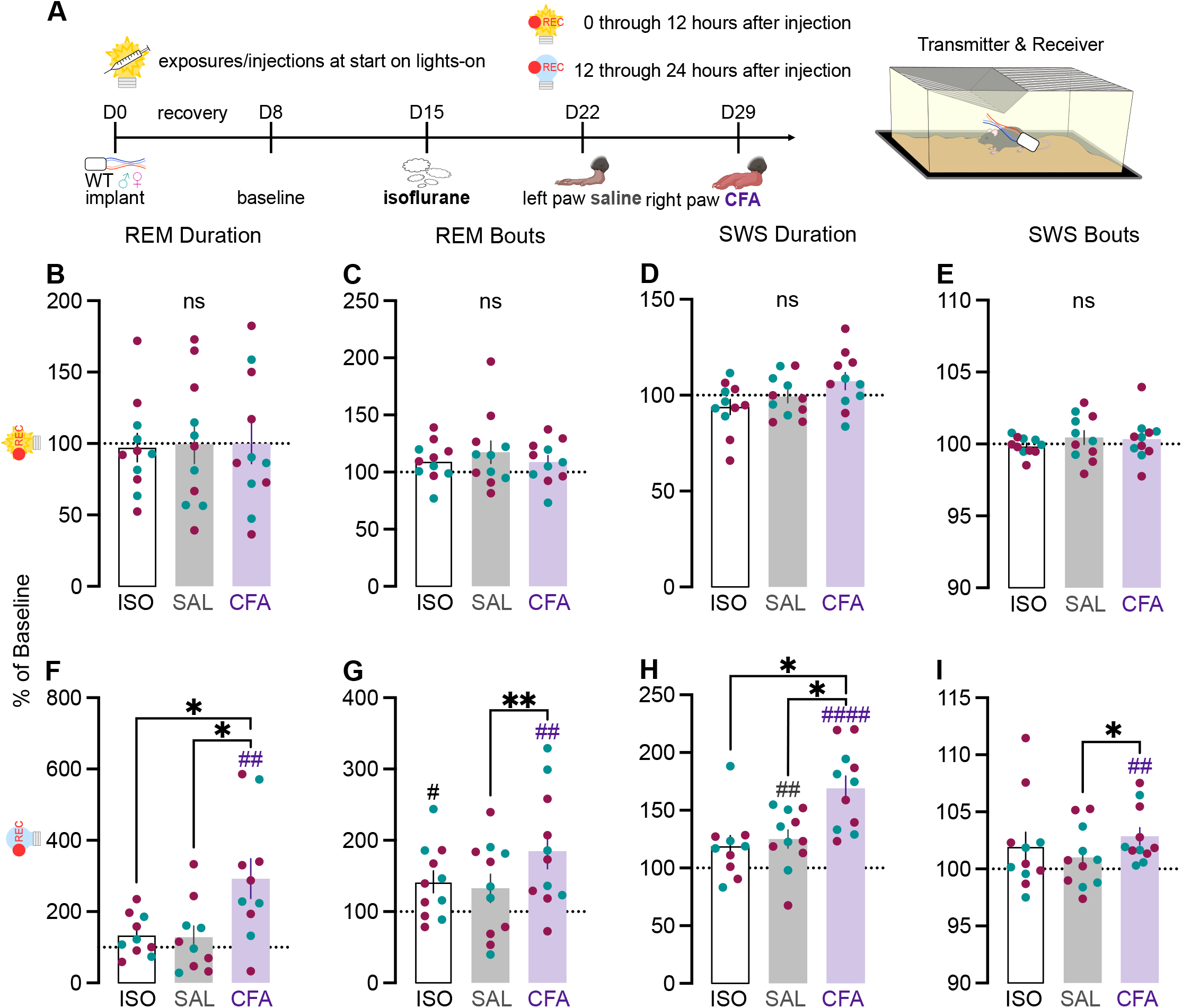
(A) Mice were implanted with wireless telemetry transmitters. EEG and EMG were recorded for 7 days of baseline, 7 days after isoflurane exposure, 7 days after saline injection, and 21 days after CFA injection. Mice were injected at the beginning of the light phase. As percent of average baseline (dotted line at 100%), (B) REM duration, (C) REM bout number, (D) SWS duration, and (E) SWS bout number during lights-on. As percent of average baseline, (F) REM duration, (G) REM bout number, (H) SWS duration, and (I) SWS bout number during lights-off. Teal points indicate males and magenta points indicate females. Ns indicates not significant. # indicates significant difference from theoretical baseline of 100% (# p<0.05, ## p<0.01, #### p<0.0001). * indicate significant group differences (* p<0.05, ** p<0.01).

We first focused our analysis on the 24 hours immediately following CFA injection. First, we confirmed that CFA reduces active wake (AW) duration. During the light phase, CFA slightly decreased AW duration (RM 1-way ANOVA F_(1.865,18.65)_=5.714, p=0.0129, Tukey’s ISO vs. CFA p=0.0346; **Fig. S1A**), and this effect was driven by females (RM 2-way ANOVA: Time F_(1.868,16.81)_=6.316, p=0.0100, Time x Sex F_(2,18)_=3.753, p=0.0434; Sidak’s Male vs. Female CFA p=0.0377; Tukey’s Female ISO vs. CFA p=0.0052, SAL vs. CFA p=0.0493; **Fig. S1A**). The AW bouts were unaffected by CFA injection (RM 1-way ANOVA F_(1.515,15.15)_=0.7527, p=0.4529; **Fig. S1B**). CFA significantly decreased locomotor activity compared to baseline, even normalized to AW duration (RM 1-way ANOVA F_(1.919,19.19)_=1.925, p=0.1741; 1-sample t-test CFA t_10_=3.383, p=0.0070; **Fig. S1C**).

During the dark phase, CFA has a stronger effect, significantly decreasing AW duration, particularly for males (RM 1-way ANOVA F_(1.740,17.40)_=10.50, p=0.0014, Tukey’s ISO vs. CFA p=0.0063, SAL vs. CFA p=0.0018; 1-sample t-test ISO t_10_=2.346, p=0.0409, SAL t_10_=2.793, p=0.0.190, CFA t_10_=9.032, p<0.0001; RM 2-way ANOVA: Time F_(1.742,15.67)_=9.381, p=0.0028, Tukey’s Male SAL vs. CFA p=0.0360; **Fig. S1D**). CFA also significantly reduced AW dark phase bouts (RM 1-way ANOVA F_(1.735,17.35)_=6.000, p=0.0129, Tukey’s SAL vs. CFA p=0.0039; 1-sample t-test ISO t_10_=3.067, p=0.0119, CFA t_10_=4.569, p=0.0010; **Fig. S1E**) specifically for males (RM 2-way ANOVA: Time F_(1.616,14.54)_=6.299, p=0.0142,; Tukey’s Male SAL vs. CFA p=0.0240; **Fig. S1E**). CFA also strongly reduced locomotor activity, particularly for females (Friedman test F=16.91, p<0.0001; Dunn’s ISO vs. CFA p=0.0042, SAL vs. CFA p=0.0004; Wilcoxon signed rank test CFA p=0.0020; 2-Way RM ANOVA Time F_(1.251,11.26)_=7.316, p=0.0160; Female SAL vs. CFA p=0.0033; **Fig. S1F**).

Next, we assessed if the reduction in wake duration is due to increases in REM, SWS, or both. During the light phase, CFA had almost no effect on REM duration (RM 1-way ANOVA F_(1.986,19.86)_=0.02847, p=0.9714; **Fig. 3B**) or bouts (RM 1-way ANOVA F_(1.427,14.27)_=0.745,p=0.4489; **Fig. 3C**). However, CFA-induced inflammation did increase SWS duration only in the females during the light phase (RM 1-way ANOVA F_(1.390,13.90)_=4.341, p=0.0457; RM 2-way ANOVA: Time F_(1.639,14.75)_=5.500, p=0.0207, Time x Sex F_(2,18)_=6.197, p=0.0090; Tukey’s Female ISO vs. CFA p=0.0133, SAL vs. CFA p=0.0302; **Fig. 3D**) without any effects on SWS bouts (RM 1-way ANOVA F_(1.960,19.60)_=0.8844, p=0.4269; **Fig. 3E**).

During the dark phase, however, CFA-induced inflammation increased both REM duration (RM 1-way ANOVA F_(1.446,13.01)_=10.72, p=0.0032, Tukey’s ISO vs. CFA p=0.0224, SAL vs. CFA p=0.0115; 1-sample t-test CFA t_9_=3.462, p=0.0071; **Fig. 3F**) and REM bouts (RM 1-way ANOVA F_(1.716,17.16)_=6.088, p=0.0126, Tukey’s SAL vs. CFA p=0.0092; 1-sample t-test ISO t_10_=2.740, p=0.0208, CFA t_10_=3.455, p=0.0062; **Fig. 3G**). REM bouts were most increased for males (RM 2-way ANOVA: Time F_(1.489,13.40)_=6.893, p=0.0130,; Tukey’s Male SAL vs. CFA p=0.0443; **Fig. 3G**).

As with REM, CFA also had much stronger effects on SWS during the dark phase. SWS duration (RM 1-way mixed effects ANOVA F_(1.976,28.66)_=8.823, p=0.0011, Tukey’s ISO vs. CFA p=0.0112, SAL vs. CFA p=0.0170; 1-sample t-test SAL t_10_=3.188, p=0.0097, CFA t_10_=6.548, p<0.0001; **Fig. 3H**) and bouts (Friedman test F=6.545, p=0.0435, Dunn’s SAL vs. CFA p=0.0315; Wilcoxon signed rank test CFA p=0.0010; **Fig. 3I**) were both significantly increased.

Next, we determined the time course of the dark-phase effects of CFA. Consistent with our piezosensor findings (**Fig. 2**), EEG and EMG recordings showed that CFA injection primarily affects sleep during the first three days post-injection (**Fig. S2**). A main effect of time was observed in wake duration (F_(4.253,34.28)_=3.907, p=0.0091; **Fig. S2A**), wake bouts (F_(4.530,36.24)_=4.094, p=0.0059; **Fig. S2D**), and REM bouts (F_(3.580,28.64)_=4.018, p=0.0126; **Fig. S2E**).

### CFA promotes sleep and suppresses wakefulness most in the dark or inactive phase

Taken together, our results indicated the CFA-induced inflammation most significantly suppressed active wakefulness, with corresponding elevations in both REM and non-REM, or slow-wave, sleep, during the dark phase when animals are typically more active. To determine whether these changes were specific to the dark phase or if CFA-induced inflammation required at least 12 hours to alter sleep, we replicated our experimental design with the exception that animals were now exposed to isoflurane (since brief anesthesia is required to prevent damage to subcutaneous leads during handling) and injected with saline and CFA at the beginning of the lights-off period.

During the dark phase, CFA significantly reduced active wakefulness duration (RM 1-way ANOVA F_(1.793,14.35)_=18.40, p=0.0001, Tukey’s ISO vs. CFA p=0.0004, SAL vs. CFA p=0.0072; 1-sample t-test

CFA t_8_=11.28, p<0.0001; **Fig. S3A**) and bouts (RM 1-way ANOVA F_(1.611,12.89)_=4.084, p=0.0493; 1-sample t-test ISO t_8_=2.413, p=0.0423, CFA t_8_=3.677, p=0.0062; **Fig. S3B**). There were no significant changes in dark phase locomotor activity (RM 1-way ANOVA F_(1.344,10.75)_=3.582, p=0.0772; **Fig. S3D**). During the light phase, CFA also significantly reduced AW duration (RM 1-way ANOVA F_(1.839,14.72)_=5.328, p=0.0200, Tukey’s ISO vs. CFA p=0.0458; 1-sample t-test CFA t_8_=3.409, p=0.0092; **Fig. S3E**) without significant effects on AW bouts (RM 1-way ANOVA F_(1.895,15.16)_=0.05948, p=0.9352;**Fig. S3F**) or locomotor activity (RM 1-way ANOVA F_(1.809,14.47)_=1.824, p=0.1979; **Fig. S3H**).

Together, these findings show that CFA injection at the beginning of the dark phase reduced active wake duration 12 to 24 hours later, during the light phase, and mice sleep even more during what is already their inactive phase. During the dark phase, however, when mice are typically more active, CFA significantly reduced active wake duration, bouts, and bout length regardless of when animals were injected.

Next, we assessed whether the timing of CFA-induced inflammation onset would differentially affect REM and SWS. REM duration during the dark phase was increased particularly for females (RM 1-way ANOVA F_(1.736,13.89)_=6.291, p=0.0136, Tukey’s ISO vs. CFA p=0.0313; 1-sample t-test CFA t_8_=3.466, p=0.0085; RM 2-way ANOVA: Time F_(1.558,10.91)_=8.791, p=0.0075, Sex F_(1,7)_=6.058, p=0.0434,; Tukey’s Female ISO vs. CFA p=0.0245; **Fig. 4B**). REM bouts (RM 1-way ANOVA F_(1.671,13.37)_=2.908, p=0.0961; 1-sample t-test ISO t_8_=2.429, p=0.0413, CFA t_8_=2.794, p=0.0234; **Fig. 4C**) were increased during the dark phase. Dark phase SWS, however, was affected more by CFA inflammation that was induced at the beginning of lights-off. During the immediate dark phase, CFA increased SWS duration, specifically in females (RM 1-way ANOVA F_(1.571,12.57)_=15.37, p=0.0007, Tukey’s ISO vs. CFA p=0.0002, SAL vs. CFA p=0.0127; 1-sample t-test CFA t_8_=9.234, p<0.0001; RM 2-way ANOVA: Time F_(1.402,9.816)_=13.83, p=0.0025; Tukey’s Female ISO vs. CFA p=0.0020; **Fig. 4D**). SWS bouts in the dark phase were also increased by CFA (RM 1-way ANOVA F_(1.683,13.47)_=5.901, p=0.0177, Tukey’s ISO vs. CFA p=0.0305, SAL vs. CFA p=0.0225; 1-sample t-test CFA t_8_=7.063, p=0.0001; **Fig. 4E**).

**Figure 4.**
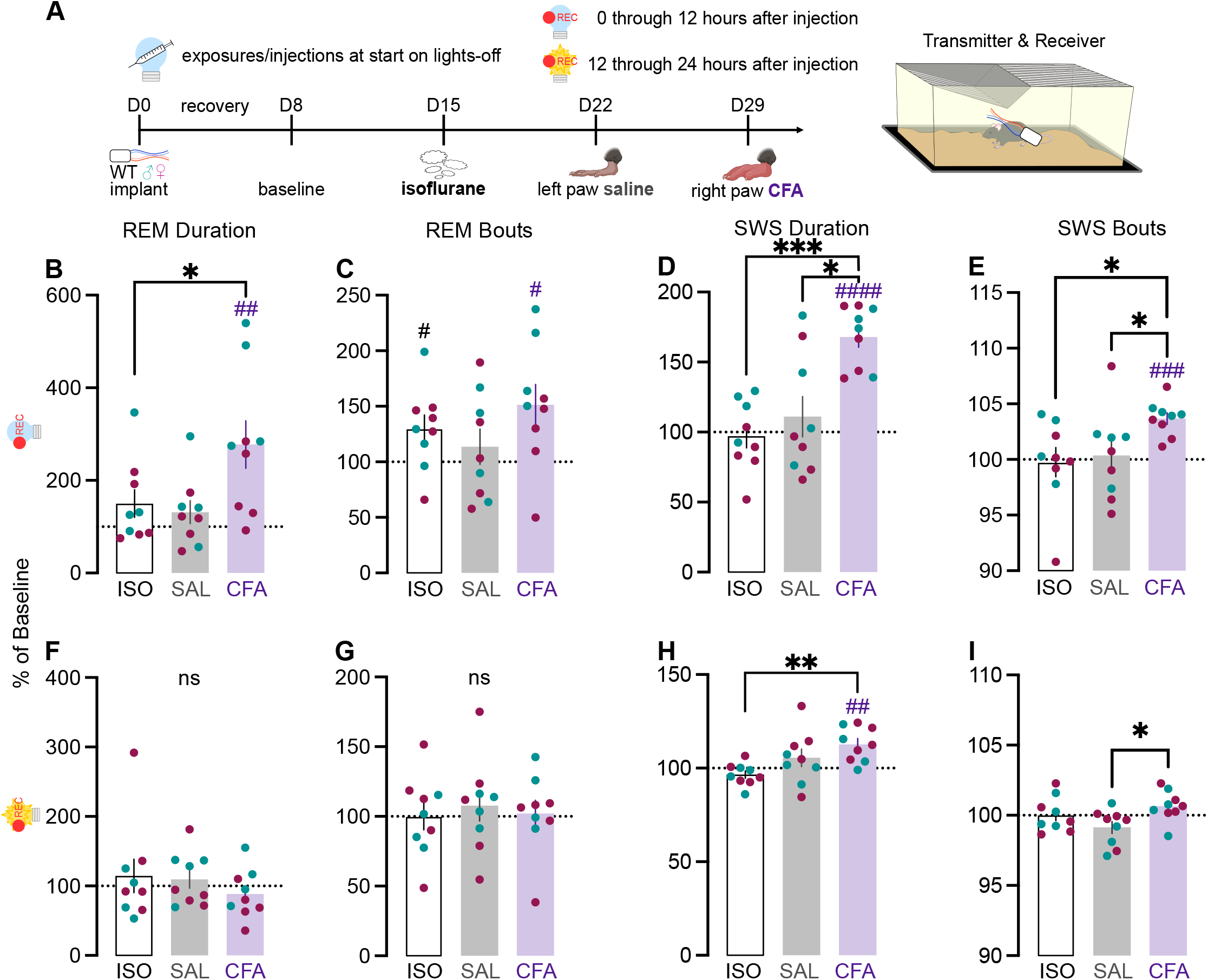
(A) Mice were implanted with wireless telemetry transmitters. EEG and EMG were recorded for 7 days of baseline, 7 days after isoflurane exposure, 7 days after saline injection, and 21 days after CFA injection. Mice were injected at the beginning of the dark phase. As percent of average baseline (dotted line at 100%), (B) REM duration, (C) REM bout number, (D) SWS duration, and (E) SWS bout number during lights-off. As percent of average baseline, (F) REM duration, (G) REM bout number, (H) SWS duration, and (I) SWS bout number during lights-on. Teal points indicate males and magenta points indicate females. Ns indicates not significant. # indicates significant difference from theoretical baseline of 100% (# p<0.05, ## p<0.01, ### p<0.001, #### p<0.0001). * indicate significant group differences (* p<0.05, ** p<0.01, *** p<0.001).

During the light phase, CFA did not affect REM duration (RM 1-way ANOVA F_(1.240,9.919)_=1.053, p=0.3478; **Fig. 4F**) or bouts (RM 1-way ANOVA F_(1.826,14.61)_=0.259, p=0.7555; **Fig. 4G**). However, CFA increased SWS duration in the light phase, specifically for females (RM 1-way ANOVA F_(1.948,15.58)_=8.956, p=0.0027, Tukey’s ISO vs. CFA p=0.0062; 1-sample t-test CFA t_8_=4.117, p=0.0034; RM 2-way ANOVA: Time F_(1.928,13.50)_=8.119, p=0.0051; Tukey’s Female ISO vs. CFA p=0.0149; **Fig. 4H**).Slow wave sleep bouts in the light phase were also increased with CFA-induced inflammation (RM 1-way ANOVA F_(1.619, 12.96)_=6.095, p=0.0175, Tukey’s SAL vs. CFA p=0.0193; **Fig. 4I**).

As seen when CFA injections were given during the light phase, EEG and EMG recordings showed that CFA injection given during the dark phase primarily affected sleep during the first two to three days post-injection (**Fig. S4**). However, CFA administered at the onset of the dark phase resulted in far more significant effects of sex, and time x sex interactions. Wake duration was significantly dynamic across three weeks of CFA (RM 2-way ANOVA F_(5.052,35.36)_=6.133, p=0.0003; **Fig. S4A**), significantly affected by sex (F_(1,7)_=6.138, p=0.0424; **Fig. S4A**) and an interaction between time and sex (F_(17,119)_=2.538, p=0.0018; **Fig. S4A**). Similarly, REM duration was significantly affected by time (RM 2-way ANOVA F_(3.094,21.66)_=12.26, p<0.0001; **Fig. S4B**), sex (F_(1,7)_=16.60, p=0.0047; **Fig. S4B**), and an interaction between time and sex (F_(17,119)_=2.356, p=0.0038; **Fig. S4B**). SWS duration was also significantly affected by time (RM 2-way ANOVA F_(4.777,33.44)_=4.762, p=0.0024; **Fig. S4C**), sex (F_(1,7)_=6.034, p=0.0437; **Fig. S4C**), and a time x sex interaction (F_(17,119)_=2.486, p=0.0022; **Fig. S4C**). Wake bouts were significantly affected by time (RM 2-way ANOVA F_(4.572,32.00)_=5.163, p=0.0018; **Fig. S4D**) and sex (F_(1,7)_=9.711, p=0.0169; **Fig. S4D**). REM bouts were affected by time only (RM 2-way ANOVA F_(4.373,30.61)_=5.381, p=0.0017; **Fig. S4E**). SWS bouts were affected by a time and sex interaction, though not by either variable independently (RM 2-way ANOVA F_(17,119)_=2.0001, p=0.0162; **Fig. S4F**). Together, these findings indicate that regardless of whether animals were injected at the beginning of lights-on (dark phase was 12 hours later) or beginning of lights-off (dark phase was immediate), CFA increased REM duration during that time. Specifically CFA-induced inflammatory pain suppresses wakefulness at the expense of both REM and SW sleep, primarily in the dark phase when mice are typically more active. This is observed regardless of whether CFA is injected at the beginning of the dark phase, or 12 hours earlier at the beginning of the light phase.

The wireless transmitters also provide subcutaneous body temperature recordings. We hypothesized that CFA-induced inflammation, and the increased sleep it causes, would also increase temperature. Surprisingly, CFA-induced inflammation significantly reduced body temperature during both lights-on, particularly for females (RM 1-way ANOVA F_(1.601,16.01)_=29.35, p<0.0001, Tukey’s ISO vs. CFA p<0.0001, SAL vs. CFA p=0.0024; 1-sample t SAL t_10_=4.477, p=0.0012, CFA t_10_=7.023, p<0.0001; RM 2-way ANOVA Time F_(1.603,14.43)_=34.25, p<0.0001, Female SAL vs. CFA p=0.0165; **Fig. S5A**) and lights-off (RM 1-way ANOVA F_(1.932,19.32)_=35.72, p<0.0001, Tukey’s ISO vs. CFA p<0.0001, SAL vs. CFA p=0.0002; 1-sample t ISO t_10_=3.495, p=0.0058, SAL t_10_=6.000, p=0.0001, CFA t_10_=10.56, p<0.0001; **Fig. S5B**).

In contrast, CFA injected at dark phase onset significantly increased body temperature during both the dark phase (RM 1-way ANOVA F_(1.715,13.72)_=20.18, p=0.0001, Tukey’s ISO vs. CFA p=0.0001, SAL vs. CFA p=0.0144; 1-sample t ISO t_8_=3.473, p=0.0084, SAL t_8_=6.292, p=0.0002, CFA t_8_=9.640, p<0.0001; **Fig. S5C**) and light phase (RM 1-way ANOVA F_(1.999,15.99)_=5.190, p=0.0183, Tukey’s p=0.0457; 1-sample t ISO t_8_=2.719, p=0.0263, SAL t_8_=5.111, p=0.0009, CFA t_8_=2.959, p=0.0182; **Fig. S5D**).

Considering that the changes in body temperature differ dramatically after light- and dark-phase CFA injections, our findings suggest that thermoregulation is differentially regulated depending on when inflammatory pain is induced.

## DISCUSSION

CFA-induced inflammatory pain profoundly alters sleep architecture in male and female mice. Injection of the hind paw, whether with CFA or saline, reduced some measures of circadian rhythmicity such as variance, period, and amplitude. CFA increased sleep duration primarily in the dark phase, while sleep bout length was decreased in the light and increased in the dark phase. Additionally, CFA reduced wake bout length, especially during the dark phase. The overall increase in sleep was due to increases in both REM and SWS. Increases in REM and SWS duration and bouts were most significant in the dark phase, regardless of whether CFA had been injected at its onset or 12 hours prior. Taken together, these results (summarized in **Table I**) indicate that inflammatory pain acutely promotes but also fragments sleep.

**Table 1.** Summary of results for each figure, including light cycle (12 hours light, 12 hours dark or constant darkness), measurement method, injection time at zeitgeber (ZT) hour, and CFA effects for all mice pooled and for sex differences.

| Figure | Light Cycle | Method | Injections | Results: CFA:<br>Pooled Sexes: | Results: CFA:<br>Sex Differences: |
| --- | --- | --- | --- | --- | --- |
| 1B – 1F | 12:12 L:D | Infrared beam breaks | 0900 hrs<br>ZT 3 | ↓ % variance<br>↓ intradaily variability<br>↓ period |  |
| 1G – 1K | 12:12 D:D | Infrared beam breaks | 0900 hrs<br>ZT 3 | ↑ relative amplitude<br>↓ period |  |
| 2 | 12:12 L:D | piezoelectric sensors | 0900 hrs<br>ZT 3 | ↑ light and dark sleep duration<br>↓ light sleep bout length<br>↓ wake bout length<br>↑ dark sleep bout length<br>↓ dark wake bout length | Males<br>↓ light wake bout length<br>Females<br>↓ dark sleep bout length |
| 3<br>S1 | 12:12 L:D | Wireless EEG | 0900 hrs<br>ZT 0 | ↓ light and dark wake duration<br>↓ dark wake bouts<br>↓ light and dark activity<br>↑ dark REM duration & bouts<br>↑ dark SWS duration & bouts | Females<br>↓ wake light duration<br>↓ dark activity<br>↑ SWS light duration<br>Males<br>↓ wake dark duration<br>↓ wake dark bouts<br>↑ REM dark bouts |
| 4<br>S2 | 12:12 L:D | Wireless EEG | 0900 hrs<br>ZT 12 | ↓ light and dark wake duration<br>↓ dark wake bouts<br>↑ dark REM duration & bouts<br>↑ light and dark SWS duration<br>↑ light SWS bouts | Females<br>↑ REM dark duration<br>↑ SWS light & dark duration |

There are few other reports on the effects of CFA on sleep in rodents. In one report, rats injected at postnatal day 1 exhibited sleep fragmentation later in adulthood, including longer sleep latency, reduced latency to REM sleep, and reduced sleep efficiency^39^. In a report describing EEG studies in adult rats, delta amplitude and SWS were only reduced with bilateral CFA injections and after 2 days of inflammation^40^. The same was also true for bilateral chronic constriction injury (CCI), a model of neuropathic pain, and skin incision^40^.

Conversely, sleep deprivation can potentiate thermal and mechanical hyperalgesia two weeks after CFA-induced inflammatory pain^19^. The same study characterized a neuronal ensemble in the nucleus accumbens, part of the limbic system and mesolimbic reward pathway, that exhibited increased activity with both nociceptive stimulation and wakefulness^19^. Activation of these neurons exacerbated nociceptive responses and reduced SWS in CCI mice but not those with sham surgeries, while inactivation had the opposite effect^19^. This ensemble primarily comprised dopamine type 1 receptor-expressing medium spiny neurons (D1-MSNs) projecting to the ventral tegmental area (VTA), which is known to regulate motivation as well as be part of the “pain matrix,” and to the preoptic area (POA) which is known to regulate sleep and wake^19^.

We have previously reported accumbal MSN regulation of sleep, in pain- and stress-naïve animals. Chemogenetic activation of D1-MSNs reduced REM duration and bouts, while inhibition increased them^14^. Activation of D2-MSNs instead increased SWS duration and bout length, indicating another way these two neuronal populations have opposing effects^14^. It remains to be elucidated whether the balance of D1- and D2-MSN activity, or their effects on sleep circuitry, are directly impacted by inflammatory pain. D1-MSNs, however, predominantly co-express the endogenous opioid neuropeptide dynorphin, which binds to kappa opioid receptors (KORs)^41^. In addition, we have previously described how dynorphin neurons are recruited by CFA-induced inflammatory pain and become hyperexcitable; blocking their effect with KOR antagonists did not affect thermal hypersensitivity but did reverse motivational deficits for natural rewards^18^. These findings further support the idea that the nucleus accumbens is an integrator for salient stimuli, memories, and expectations for behavioral selection^42^, and that this region is a key element of brain circuitry that is modified in response to experience including stress^43^. According to this working model, the stress of pain imparts behavioral avoidance that prevents further injury or damage, and hopefully promotes recovery; but as pain becomes chronic, this avoidance becomes maladaptive as patients withdraw socially, become less mobile, and experience greater fatigue^42^.

The conceptualization of pain as a stressor is not novel, particularly given the tremendous overlap in circuitry. This overlap provided a strong justification for the present studies. Pain is highly comorbid with stress-associated disorders where around 30% of patients in pain also meet diagnostic criteria for major depressive disorder (MDD)^9,10^. Our lab has previously reported chronic social defeat stress, a widely used mouse model for chronic stress given its sensitivity to chronic antidepressants and its propensity to stratify mice into resilient or susceptible to the stressor, increased both REM and SWS duration over 10 days of defeats^11^. Furthermore, the increase in REM was reduced by a systemically administered long-acting KOR antagonist, again indicating common neural substrates for sleep, stress, and pain^11^. Future work involving newly developed short-acting KOR antagonists may enable more precise characterization of the roles of KORs in the acute and chronic effects of pain and associated stress-related states.

Future studies are needed to determine the mechanisms by which inflammatory pain promotes sleep, which will enable the development of improved therapeutics. Clinically, inflammatory pain includes fibromyalgia and rheumatoid arthritis (RA). CFA-induced inflammatory pain closely emulates the latter, with some studies identifying rheumatoid factor in the joints of CFA-injected rodents^44,45^. RA patients report both longer sleep duration and more frequent daytime naps, and their fatigue is highly correlated with individual ratings of pain and stiffness^46^. Inflammation and sleep disturbances, particularly in RA, can have a reciprocal or feedforward relationship, and even make pain sensitivity worse^47^. From a translational perspective, RA develops slowly but progressively over time, and thus targeting a discrete time point for preventing increased sleep is not feasible. However, as RA becomes chronic and its symptoms worse, fatigue and drowsiness may severely impair the quality of life for patients as much as their reduced mobility and pain. In rodent studies, increased sleep may be the direct result of fatigue and drowsiness rather than a specific promotion of sleep or suppression of wake, as rodents have no need to keep themselves awake, particularly in a laboratory setting. The present studies primarily focus on the first week after the induction of inflammatory pain, and future priorities include parsing whether these described effects on sleep architecture are due to the injection, inflammation, pain, or all three.

Another area that remains to be investigated is whether baseline or subsequent individual differences in sleep traits and inflammatory processes contribute to increased sleep in inflammatory pain. This type of heterogeneity has been well-studied in preclinical stress research, particularly in the CSDS model^48^. Are there any baseline sleep traits, whether that be duration or latency to certain vigilance states or spectral power, which can predict either the hyperalgesia or increased sleep of CFA-induced inflammatory pain? If so, would modulating these traits alter the ultimate hyperalgesia or sleep profile of rodents with inflammation, or inflammatory pain? The search for biomarkers, whether indicative of a symptom or its effective treatment, will be aided by the methods this study employed, namely, continuous, objective, and translational phenotyping. Moreover, as novel therapeutics are developed, leveraging sleep telemetry as a preclinical research endpoint will be vital to characterizing not only how new drugs affect disease symptoms, but also overall quality of life.

In summary, our findings that CFA increases and fragments sleep during the rodent active phase provide further characterization to this widely used pain model, and open the door to many future studies for identifying neuronal ensembles and neural circuits that commonly mediate both nociception and sleep behaviors.

## DATA AVAILABILITY

Authors can confirm that all relevant data are included in the paper and its Supplementary Information files. All data generated in this study are provided in the Supplementary Information/ Source Data file.

## AUTHOR CONTRIBUTIONS

Conceptualization: D.J.B., K.M.I., J.A.M., and W.A.C.

Methodology: D.J.B., K.M.I., A.S., E.S.M., J.A.M., and W.A.C.

Formal analysis: D.J.B., K.M.I., and A.S.

Investigation: D.J.B., K.M.I., and A.G.H.

Writing – Original Draft: D.J.B.

Writing – Reviewing & Editing: K.M.I., A.S., E.S.M., J.A.M., and W.A.C.

Funding Acquisition: D.J.B., J.A.M., and W.A.C.

Resources: E.S.M., J.A.M., and W.A.C.

Supervision: J.A.M., and W.A.C.

