## Supplemental for "Inflammatory pain in mice induces light cycle-dependent effects on sleep architecture"

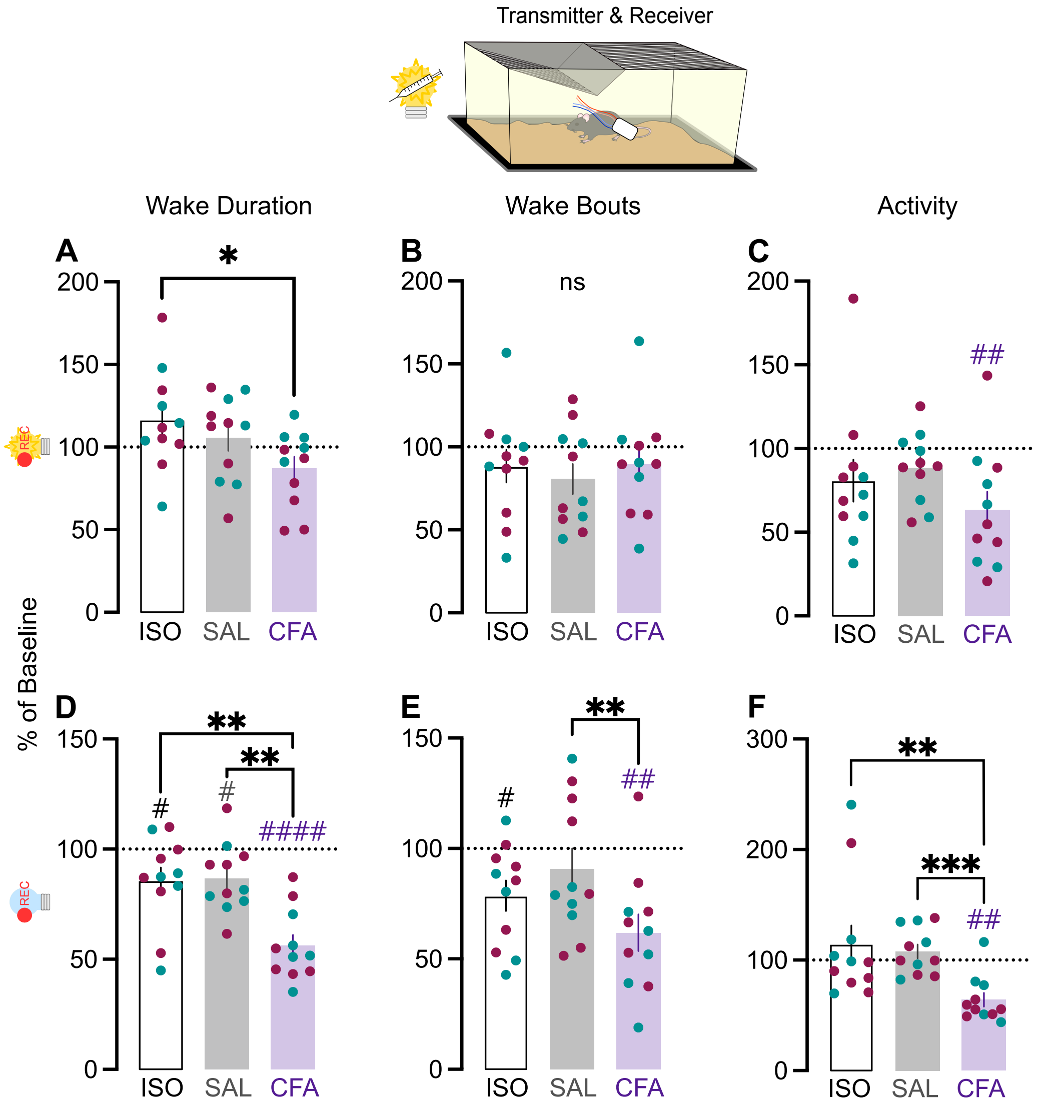


**Figure S1**. (A) Mice were implanted with wireless telemetry transmitters. EEG and EMG were recorded for 7 days of baseline, 7 days after isoflurane exposure, 7 days after saline injection, and 21 days after CFA injection. Mice were injected at the beginning of the light phase. As percent of average baseline (dotted line at 100%), (B) wake duration, (C) wake bout number, and (D) locomotor activity during lights-on. As percent of average baseline (D) wake duration, (E) wake bout number, and (F) locomotor activity during lights-off. Teal points indicate males and magenta points indicate females. Ns indicates not significant. # indicates significant difference from theoretical baseline of 100% (# p<0.05, ## p<0.01, #### p<0.0001). * indicate significant group differences (* p<0.05, ** p<0.01, ***p<0.001).

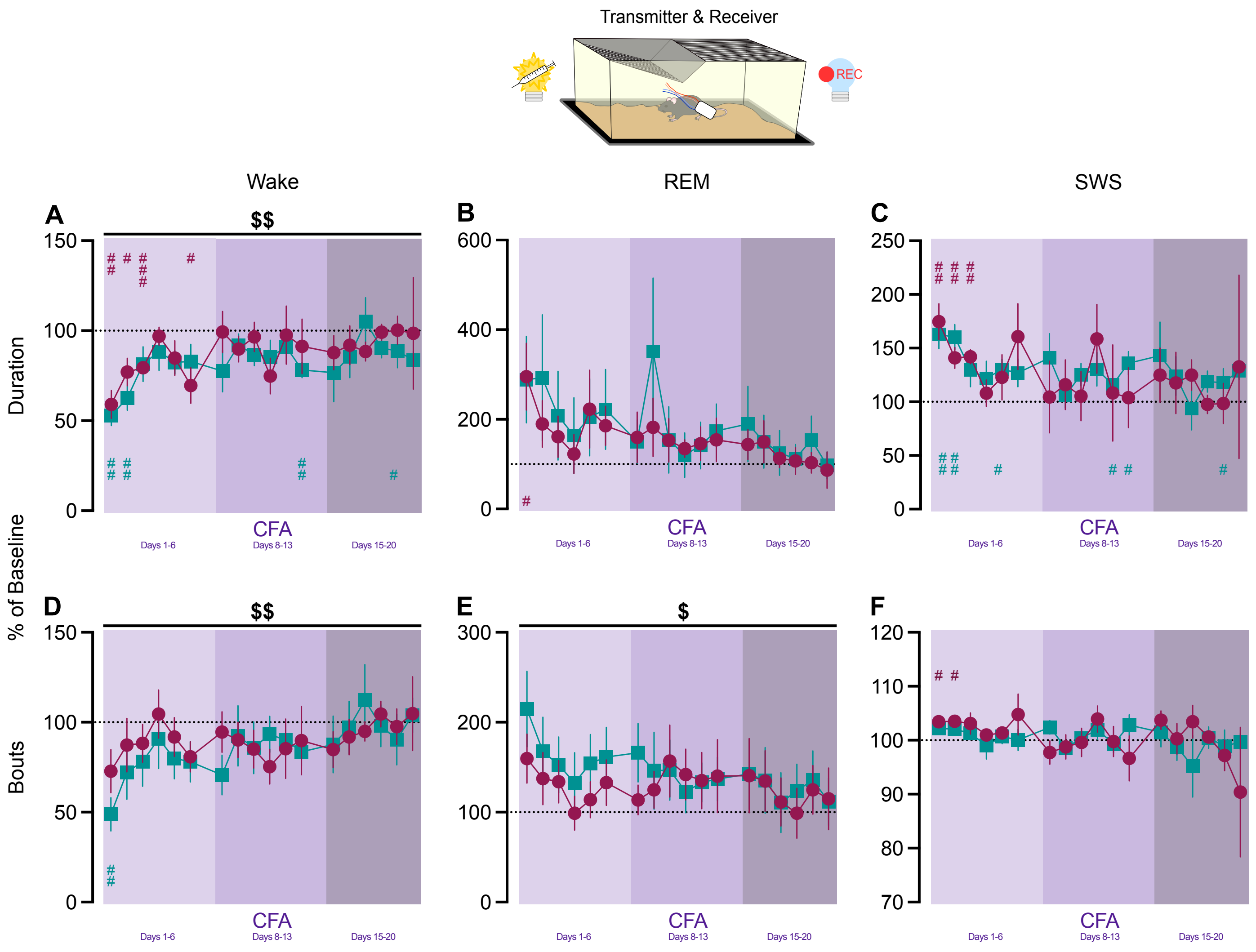


**Figure S2**. Mice were implanted with wireless telemetry transmitters. EEG and EMG were recorded for 7 days of baseline, 7 days after isoflurane exposure, 7 days after saline injection, and 21 days after CFA injection. Mice were injected at the beginning of the light phase. As percent of average baseline (dotted line at 100%), (A) wake duration, (B) REM duration, and (C) SWS duration during lights-on. As percent of average baseline (D) wake bouts, (E) REM bouts, and (F) SWS bouts during lights-off. Teal points indicate males and magenta points indicate females. # indicates significant difference from theoretical baseline of 100% (# $p<0.05$, ## $p<0.01$, ### $p<0.001$). $ indicates significant main effect of time ($ $p<0.05$, $$ $p<0.01$).

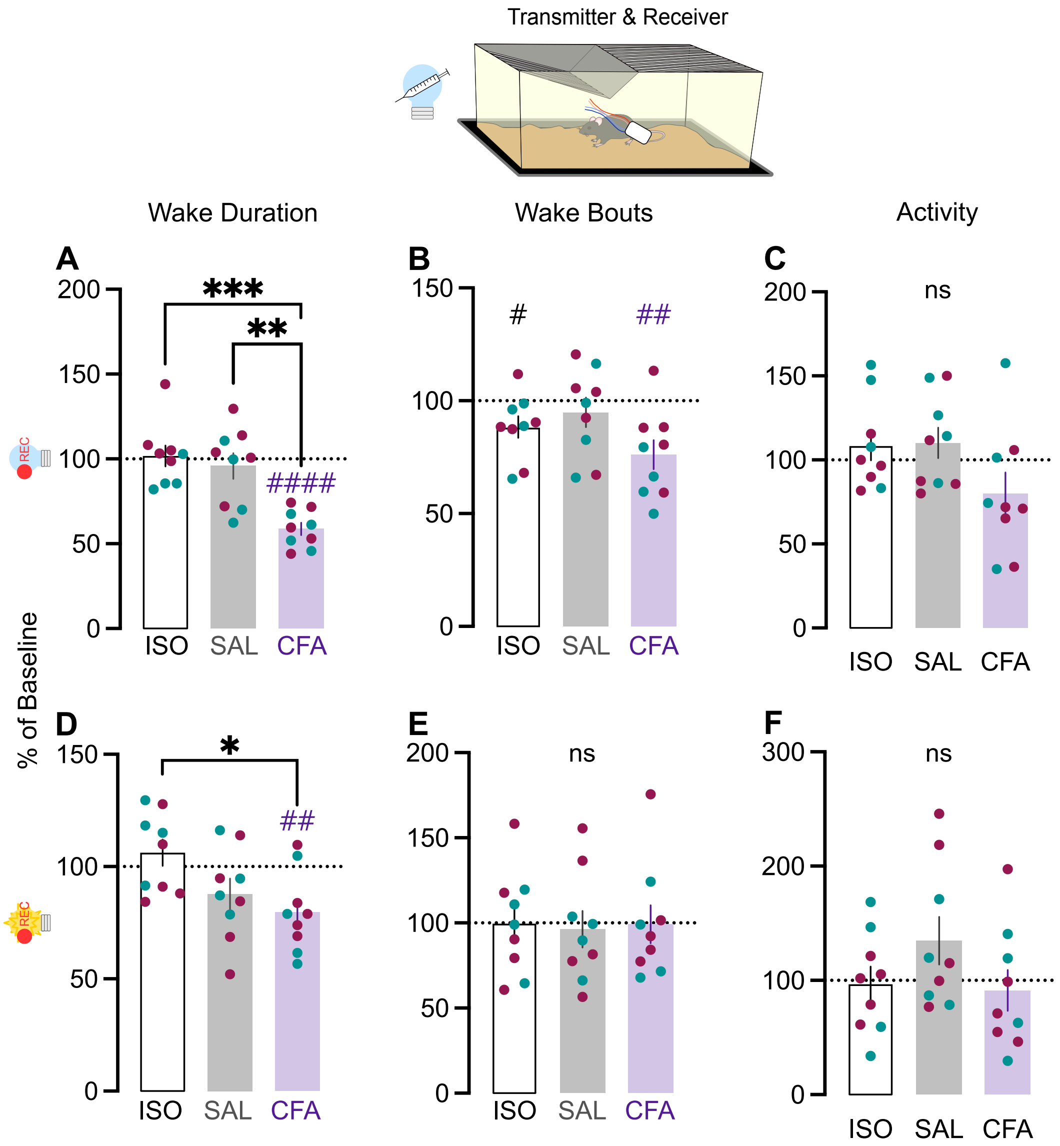


**Figure S3**. (A) Mice were implanted with wireless telemetry transmitters. EEG and EMG were recorded for 7 days of baseline, 7 days after isoflurane exposure, 7 days after saline injection, and 21 days after CFA injection. Mice were injected at the beginning of the dark phase. As percent of average baseline (dotted line at 100%), (B) wake duration, (C) wake bout number, and (D) locomotor activity during lights-off. As percent of average baseline (D) wake duration, (E) wake bout number, and (F) locomotor activity during lights-on. Teal points indicate males and magenta points indicate females. Ns indicates not significant. # indicates significant difference from theoretical baseline of 100% (# p<0.05, ## p<0.01, #### p<0.0001). * indicate significant group differences (* p<0.05, ** p<0.01, ***p<0.001).

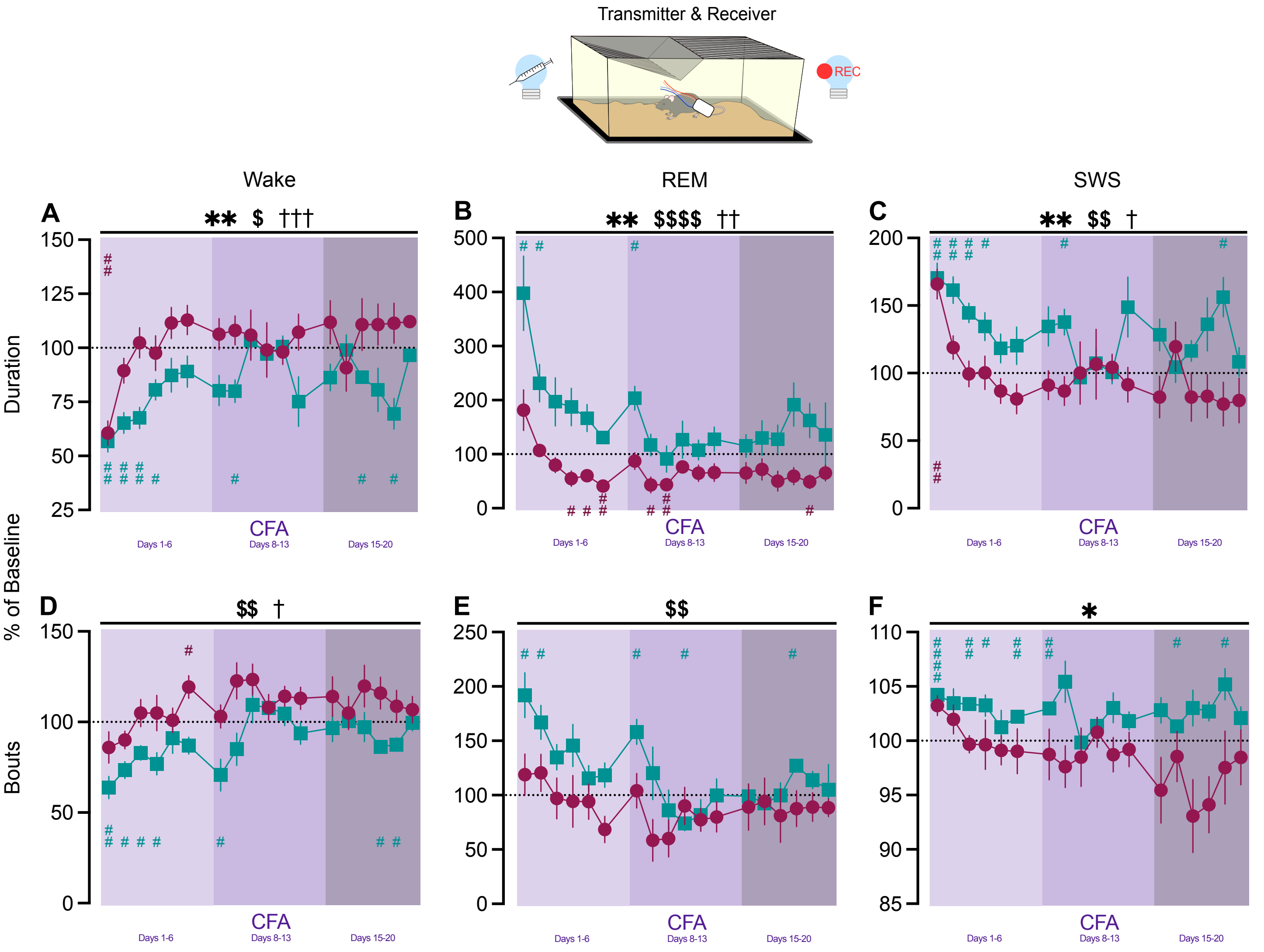


**Figure S4**. Mice were implanted with wireless telemetry transmitters. EEG and EMG were recorded for 7 days of baseline, 7 days after isoflurane exposure, 7 days after saline injection, and 21 days after CFA injection. Mice were injected at the beginning of the dark phase. As percent of average baseline (dotted line at 100%), (A) wake duration, (B) REM duration, and (C) SWS duration during lights-on. As percent of average baseline (D) wake bouts, (E) REM bouts, and (F) SWS bouts during lights-off. Teal points indicate males and magenta points indicate females. # indicates significant difference from theoretical baseline of 100% (# p<0.05, ## p<0.01, #### p<0.0001). * indicates significant main effect of time and sex interaction (* p<0.05, ** p<0.01). $ indicates significant main effect of time ($ p<0.05, $$ p<0.01, $$$$ p<0.0001). † indicates significant effect of sex († p<0.05, †† p<0.01, ††† p<0.001).

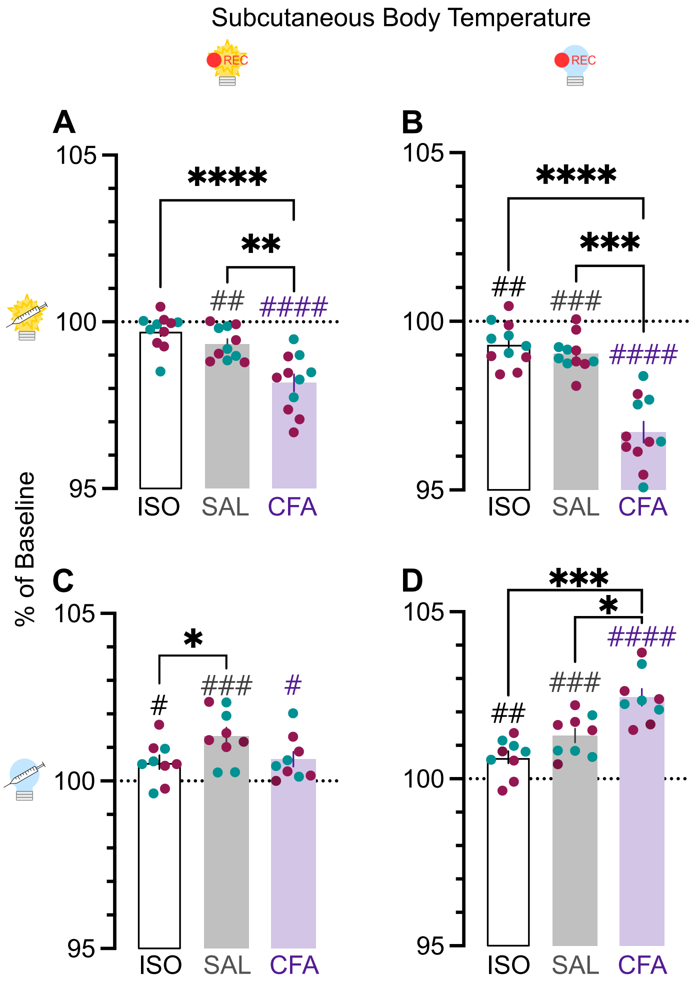


**Figure S5**. As percent of average baseline (dotted line at 100%), (A) temperature during lights-on for animals injected at the beginning of the light phase, (B) temperature during lights-off for animals injected at the beginning of the light phase, (C) temperature during lights-on for animals injected at the beginning of the dark phase, (D) temperature during lights-off for animals injected at the beginning of the dark phase. Teal points indicate males and magenta points indicate females. # indicates significant difference from theoretical baseline of 100% (# $p<0.05$, ## $p<0.01$, ### $p<0.001$, #### $p<0.0001$). * indicate significant group differences (* $p<0.05$, ** $p<0.01$, *** $p<0.001$, **** $p<0.0001$).
